# Sourmash Branchwater Enables Lightweight Petabyte-Scale Sequence Search

**DOI:** 10.1101/2022.11.02.514947

**Authors:** Luiz Irber, N. Tessa Pierce-Ward, C. Titus Brown

## Abstract

We introduce branchwater, a flexible and fast petabase-scale search for the 767,000 public metagenomes presently in the NCBI Sequence Read Archive. Our search is based on the FracMinHash k-mer sketching technique and can search all public metagenomes with 1000 query genomes in approximately 36 hours using 50 GB of RAM and 32 threads. Branchwater is a Rust-based multithreading front-end built on top of the sourmash library. We provide biological use cases, examine performance, and discuss design and performance considerations.

## Introduction

Substantial growth in publicly available nucleotide sequencing data (DNA and RNA) has occurred over the last decade, driven by decreases in sequencing costs. In particular the Sequence Read Archive now has over 9 million entries containing 12 PB of data [1]. Shotgun metagenomes, generated by random sequencing of mixtures of microbes sampled from a microbiome are a particularly interesting resource stored in the SRA.

Shotgun metagenome data sets are often large (100s of MBs to 10s of GB) and can be highly complex, with environmental samples containing genomic data that can be attributed to thousands or more species. In the past decade, hundreds of thousands of new bacterial and archaeal genomes have been isolated from public metagenomes, and several entirely new branches of life have been discovered purely through analysis of public data [2,3,4].

Beyond their initial use, these data sets form an incredibly rich resource for contextualizing novel sequencing data and for synthesis research on a myriad of large-scale genomic questions ranging from basic evolutionary processes to disease associations and pathogenicity tracking (Table 1). However, comprehensive discovery of relevant data sets is challenging. Metadata for these data sets is typically geared towards the submitting researcher’s study questions and major findings, and moreover cannot possibly describe the full contents of the data. Furthermore, metadata provided at the time of submission can be incomplete or inconsistent, rendering systematic data set discovery intractable.

**Table 1:** Biological use cases for petabase scale sequence search of metagenomes.

| Use case | Description | Enabled by branchwater |
| --- | --- | --- |
| Biogeography of genomes | Describe and characterize biogeographical distribution of species; identify sampling locations | Yes |
| Outbreak tracking | Trace pathogen spread via public data | Yes, for genomes > 10kb |
| Pangenome expansion | Expand and explore composition of strains, species and genus level pangenomes (including SAGs and MAGs) | Yes |
| SRA metadata reannotation | Content-based validation and reannotation of SRA metagenomes | Yes |
| Private database access and search | Provide privacy-enabled search of large, access-restricted databases | Yes |
| Post-processing and cleaning MAGs | Evaluating contig-level presence and abundance across multiple metagenomes | Yes, for contigs > 10kb |
| Exploring breadth of plasmids etc. | Evaluating range and prevalence of laterally transferred genomic elements | Yes, for plasmids > 10kb |
| Exploring host range of species for regulatory evaluation |  | Yes, for genomes > 10kb |
| Search for small viruses |  | No |
| Searching for specific functional genes |  | No |
| Detecting novel classes and orders |  | No |

Content-based search is a promising alternative strategy for finding data sets in archives. By searching with genomic content of interest, content-based search can recover datasets containing relevant species or genes of interest regardless of their associated metadata. However, search of unassembled sequence is critical to ensure unbiased and comprehensive recovery of relevant datasets. Assembly techniques are designed to produce consensus reference sequences useful for consistent comparisons across genotypes, often collapsing sequence variation in the process [5,6]. In addition, reassembly and reanalysis of existing data using different parameters or newer methods often yields different results. Content-based search of unassembled metagenomes can bypass these issues and facilitate consistent downstream analysis across data sets that may have been initially generated to answer a range of disparate biological questions, and been first analyzed over a range of years and with myriad techniques.

A number of approaches have been developed to enable content-based search of single-organism genomic and RNAseq data. Methods that enable rapid, large-scale search across hundreds of thousands of data sets typically leverage biological sketching techniques and probabilistic data structures to reduce the effective search space [7,8,9]. However, these approaches do not readily translate to datasets with unknown levels of sequence diversity, the defining feature of metagenomic datasets.

Recent Serratus used extensive search across public datasets to recover 880,000 RNA-dependent RNA polymerase-containing sequences, and discovered over 131,000 novel RNA viruses [10]. This search was comprehensive but also time-consuming and costly, and still out of reach for unfunded exploratory research. Other approaches such as searchsra.org [11] and metagraph [12] are promising but are not yet capable of searching all public data.

Here, we introduce Branchwater, a petabase-scale querying system that uses containment searches based on FracMinHash sketching to search all public metagenome data sets in the SRA in 24-36 hours on commodity hardware with 1 −1000 query genomes. Branchwater uses the Rust library underlying the sourmash implementation of FracMinHash to execute massively parallel searches of a presketched digest of the SRA [13,14].

The availability of relatively lightweight content-based search of SRA metagenomes helps address many of the biological use cases in Table 1 (see 3rd column). Some of these use cases have already been explored with Branchwater: Viehweger et al. (2021) [15] used Branchwater to discover a metagenomic sample containing *Klebsiella pneumonia* that was subsequently included in an outbreak analysis, and Lumian et al. (2022) [16] conducted a biogeographical study on five newly generated cyanobacterial genomes from Antarctic samples.

## Background: FracMinHash and sourmash

FracMinHash is a bottom-sketch version of ModHash that supports accurate estimation of overlap and containment between two sequencing sets [14]. In brief, FracMinHash is a lossy compression approach that represents data sets using a “fractional” sketch containing 1/S of the original k-mers. FracMinHash sketches support estimation of overlap, bidirectional containment, and Jaccard similarity between two data sets. Unlike other common sketching techniques such as MinHash [17] and HyperLogLog [18], FracMinHash supports these operations between two data sets of different sizes, which is important for metagenomic search; and unlike mash screen and CMash, FracMinHash does not require the original data sets [19,20]. In exchange, FracMinHash sketches are essentially unbounded in size, since they can grow to include up to ***H/S*** elements for a hash space ***H*** in size.

The open-source sourmash software provides a mature and well-documented command-line interface to FracMinHash, along with Python and Rust APIs for loading and using FracMinHash sketches [13,21]. The Python layer provides a larger number of user experience conveniences on top of the performant Rust layer. However, despite the thread safety of the underlying Rust code, the CLI and Python library still operate in single-threaded mode, which limits the utility of sourmash for very large scale operations. Refactoring the sourmash CLI and Python libraries to take advantage of thread safety is a substantial and ongoing effort; for petabase scale search, we chose to develop a dedicated CLI in Rust in the interim.

There are several features of FracMinHash and sourmash that limit their utility for specific use cases. In particular, the default ***S*** = **1000** parameter used in sourmash does not work well for comparing or detecting genomes smaller than 10kb in size. Nor can highly divergent genomes be found; based on k-mer containment to ANI conversion [22], we find that sourmash defaults work well for finding matches to genomes within about 90% ANI of the query, but not necessarily further. Finally, FracMinHash was developed for shotgun data sets and different parameters would be required for targeted sequencing data such as amplicon data sets. Some of these limitations are intrinsic to FracMinHash, and others may be overcome in the future by parameter tuning and further research.

### Petabase scale search represents a specific technical challenge to sourmash

The primary design focus for the sourmash CLI has been on searching and comparing many microbial genome-sized sketches, where for typical parameters there are between 1000 and 10,000 hashes in each sketch. The software provides a variety of in-memory and on-disk data structures for organizing sketches in this size range and can search hundreds of thousands of genome sketches with a single query in minutes in a single thread on an SSD laptop; more complex algorithms such as the min-set-cov approach described in [14] can take a few hours but are still acceptably performant on most real-world data.

Branchwater faces very different challenges in searching large collections of *metagenomes*. Many of these data sets are extremely large, slow to read from disk, and individually require substantial memory to load. Where multiple queries are used to search each metagenome, quadratic search costs will also be incurred.

One solution we explored was a scatter-gather approach based on a cluster-aware workflow engine (in this case, snakemake [23]). The overhead on workflow coordination and executing shell commands was prohibitive for our initial implementation, so we pursued a purpose-built multithreaded solution instead.

## Methods

### Sketching the Sequence Read Archive

We determined the accessions of all publicly available shotgun metagenomic via the query string `“METAGENOMIC”[Source] NOT amplicon[All Fields]` at the NCBI Sequence Read Archive Web site, https://www.ncbi.nlm.nih.gov/sra. We then downloaded all runs for all accessions and streamed them into sourmash sketch dna with parameters `-p k=21,31,51,scaled=1000,abund`. The output sourmash signature files were saved as individual gzipped JSON files (each containing 3 sketches), one file for each input run.

The resulting catalog contains 767,277 metagenome data sets as of March 2022, with the largest category annotated as human-associated microbiomes (Table 2). The size of all sketches together is 7.5 TB, containing approximately 375 billion hashes per k-mer size, representing 375 trillion k-mers. There are 2.20 billion distinct hashes in this collection, representing approximately 2.20 trillion distinct k-mers.

**Table 2:** The 10 largest categories of metagenome data set types in the Sequence Read Archive, as of March 2022.

| "Scientific Name" provided by submitter | distinct data sets |
| --- | --- |
| human gut metagenome | 162187 |
| metagenome | 57048 |
| gut metagenome | 47244 |
| human metagenome | 36438 |
| soil metagenome | 35323 |
| mouse gut metagenome | 26482 |
| human skin metagenome | 25700 |
| Homo sapiens | 21020 |
| marine metagenome | 14400 |
| human oral metagenome | 14235 |

The average sketch file size is 9.7 MB, and the median file size is 570kb. The largest 10,000 data sets comprise 30% of the total sketch sizes.

### Implementation of multithreaded search

The branchwater program is built in Rust on top of the sourmash library for loading and comparing sketches. It implements the following steps:

1. Loads all query sketches into memory from a list of files.
2. Loads the list of filenames containing subject sketches to search.
3. In a Rust closure function executed in parallel for each subject sketch filename,

a. loads the subject sketch from the file;
b. for each query, determines the estimated overlap between query and subject;
c. reports overlaps above a user-specified threshold.
d. releases all per-metagenome resources

Downsampling of sketches to higher scaled values is performed dynamically, after load (if requested). Results are reported back to a separate “writer” thread via a threadsafe multi-producer, single-consumer FIFO queue. We use the rayon `par_iter` function to execute the closures in parallel.

This approach leverages the core features of sourmash to efficiently keep queries in memory and batch-process metagenome sketches without storing them all in memory. The approach also takes advantage of the effective immutability of queries, which can be shared without data races by multiple processing threads.

### Executing branchwater at the command line

branchwater takes in search parameters as well as two text files, one containing a list of query file paths and one containing a list of subject file paths. Upon execution, it reports the number of query sketches loaded and the number of subject file paths found, and then begins the search. It progressively reports the number of sketches searched in blocks of 10000, and outputs matches to a CSV File.

We typically run branchwater in a snakemake workflow, which manages environment variables and input/output files.

## Results

### Example Branchwater search results

To showcase the utility of Branchwater search, we searched a gut bacterium mixture against all indexed SRA metagenomes using a k-mer size of 31. Branchwater search is exhaustive: every query is searched against every search metagenome. Reporting resources are minimal: each match is reported as a single line in a CSV containing the query and match identities and the k-mer containment of the query in the match. As a result, we use a low default threshold, 0.01, which means that any metagenome that contains more than 1 % of the k-mers in any query is reported. Branchwater search with this default threshold returned 192,699 metagenomes. We then filtered these results for metagenomes that contained at least 20% of query k-mers, retaining 66,705 metagenomes for further analysis. The summarized SRA metadata (Table 3) provides some insight into the types of metagenomes recovered: the majority were annotated as “human gut metagenome” or “gut metagenome”, with a smaller number of other gut-related categories, including “wastewater metagenome” and “mouse gut metagenome”. Branchwater also reported 167 “sediment metagenome” samples, but upon further investigation, many of these originated from a transect study investigating the presence of microbes with distance from wastewater treatment.

**Table 3:**
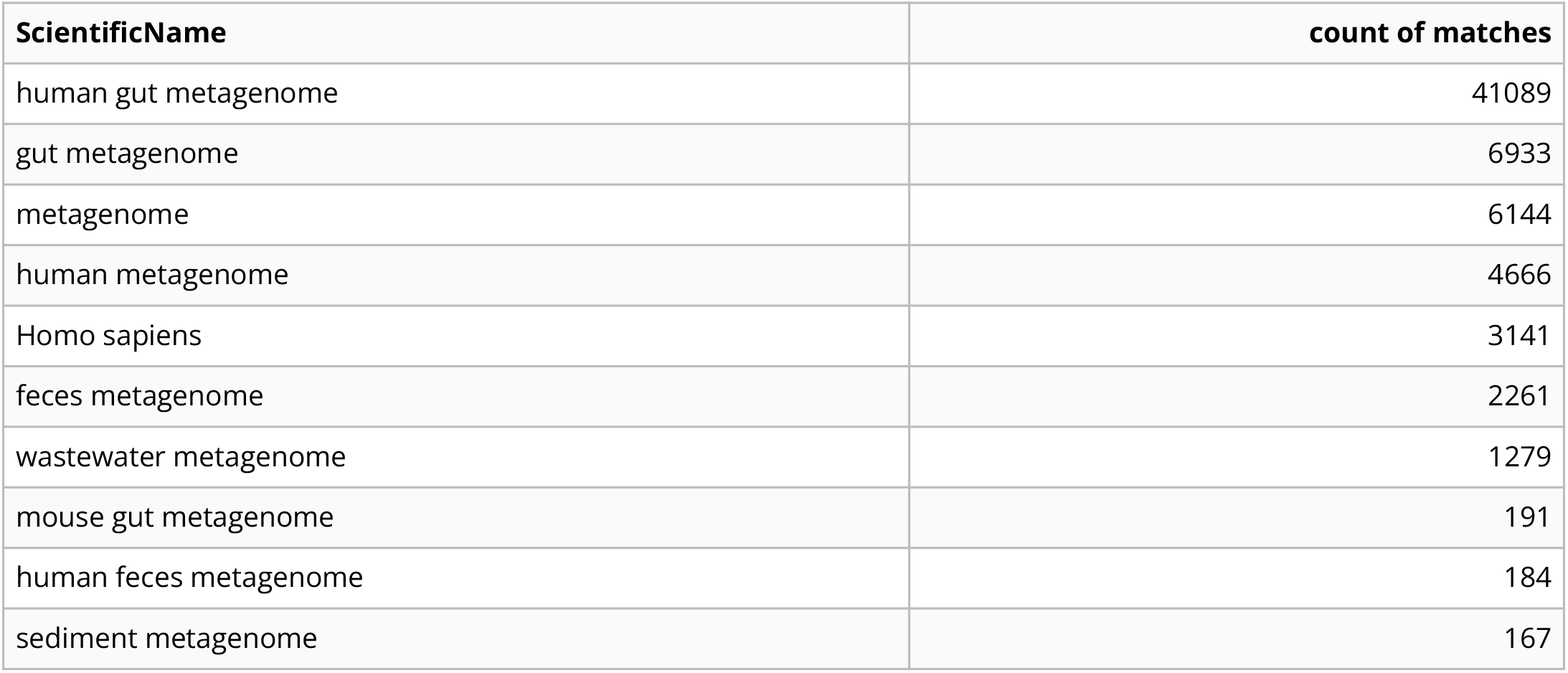
Output summary of ScientificNames from metagenome annotations in the Sequence Read Archive for a gut bacterium search mixture. We have provided a simple script that imports the SRA metadata and summarizes the Branchwater results at the provided threshold.

| ScientificName | count of matches |
| --- | --- |
| human gut metagenome | 41089 |
| gut metagenome | 6933 |
| metagenome | 6144 |
| human metagenome | 4666 |
| Homo sapiens | 3141 |
| feces metagenome | 2261 |
| wastewater metagenome | 1279 |
| mouse gut metagenome | 191 |
| human feces metagenome | 184 |
| sediment metagenome | 167 |

### Performance and scaling analysis

In Tables 4 and 5 we show performance metrics for branchwater. Table 4 contains average measurements and standard deviations for time, memory, and I/O, showing that branchwater is I/O and memory intensive. Table 5 compares the runtimes and memory usage for a search the entire catalog (normalized to 10k samples) against both the random subsets in Table 4 and the largest 10k sketches in the database. The increased memory usage of the catalog and especially the 10k largest metagenomes suggests that a relatively small number of metagenomes contributes the bulk of both time and memory usage to a full catalog search of branchwater.

**Table 4:**
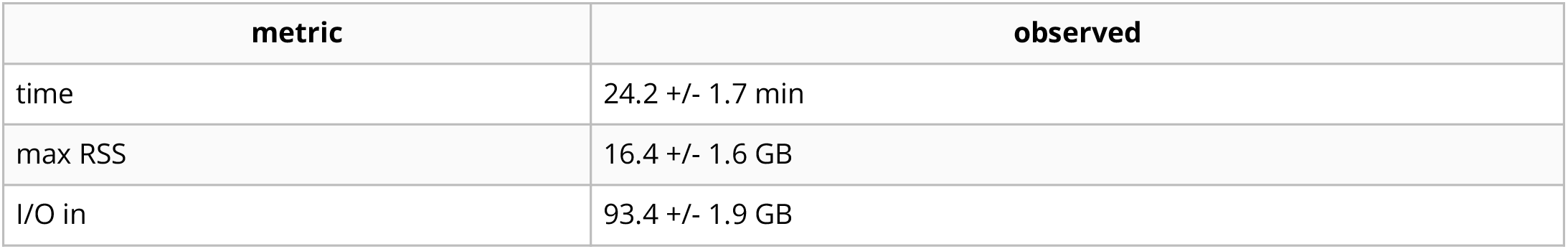
Time, memory, and I/O input for 5 runs of 1000 queries against 10,000 metagenomes. Queries were randomly selected from 318k genomes in GTDB rs207. Metagenomes were randomly selected from the full catalog of 767k.

| metric | observed |
| --- | --- |
| time | 24.2 +/- 1.7 min |
| max RSS | 16.4 +/- 1.6 GB |
| I/O in | 93.4 +/- 1.9 GB |

**Table 5:** A comparison of database searches for the entire catalog of 767k metagenomes, with time normalized to 10k, the average of the 10k subsets from Table 4, and the 10k largest metagenomes.

| database | time | max RSS |
| --- | --- | --- |
| entire catalog | 27.2m* | 52.5 GB |
| random 10k subset | 24.2m | 16.4 GB |
| 10k largest metagenomes | 594.0 m | 60.0 GB |

**Figure 1:**
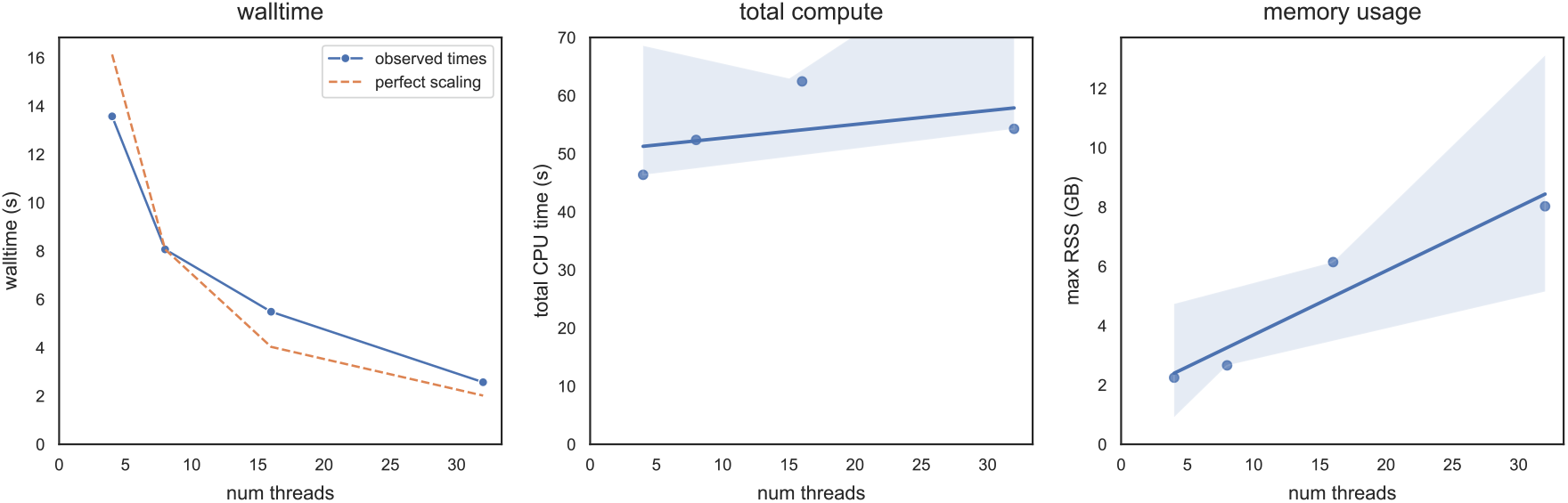
branchwater scales well with number of threads. Processing time drops linearly with number of threads, while total compute stays approximately the same and memory usage increases linearly with number of threads as each thread loads a subject to search.

**Figure 2:**
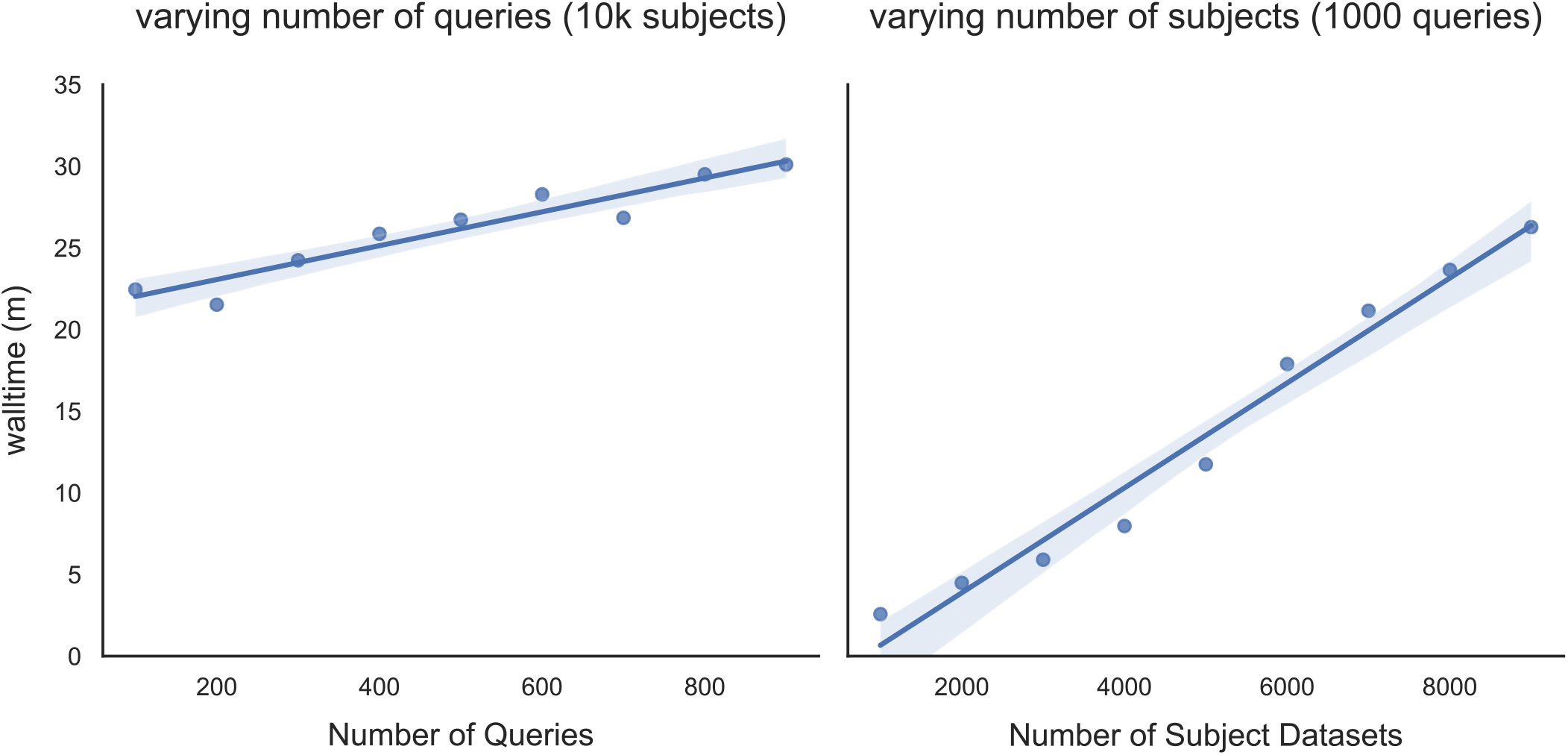
branchwater scales linearly with number of queries and subjects, but number of subjects dominates runtime. Processing time increases slowly with number of query genomes used to search, because they are held in memory and fast to compare. Processing time increases quickly with number of subject metagenomes being searched, because they are large and slow to load and search.

See https://github.com/dib-lab/2022-branchwater-benchmarking for number processing and figure generating notebooks.

### Post-search validation

The sourmash CLI can be used to explore k-mer matches for individual data sets. This does not validate the matches beyond confirming the containment numbers, although sourmash provides additional information (e.g. estimated abundances) on top of the minimal information provided by branchwater. FracMinHash generally and Branchwater specifically have been validated bioinformatically with read mapping (see [14,16]). This is discussed further below.

## Discussion

Enabling content-based search of very large collections of sequencing data is an open problem, and approaches that work for smaller collections rarely scale well, even for current database sizes. New methods that take advantage of specific particularities of the query and desired answer can help bridge the gap between more general methods by allowing filtering large databases, resulting in more manageable subsets that can be used efficiently with current methods.

K-mer search via sourmash branchwater is a lightweight, scalable approach to content-based search of petabase-scale collections of sequencing data. Branchwater is designed to recover data sets with high nucleotide similarity to the query sequence (>90% ANI) across the Sequence Read Archive using computational resources available to many researchers. As this approach enables content-based search across petabases of sequence data, we anticipate it will be most useful for filtering large databases to generate a manageable subset of relevant data sets that can be analyzed in detail using other tools.

Despite its utility, branchwater search has limitations, some of which are intrinsic to the approach, and some of which can be overcome with improved database storage and search design. First, branchwater search relies upon exact matching of long nucleotide k-mers, which work best for sequences with high sequence identity (90%+ ANI), particularly when combined with thresholding used for fast search. Approaches using alignment-based techniques may be more useful for detecting similarity across larger evolutionary distances [10]. Lumian et al. (2022) validated branchwater results by downloading matching Illumina metagenomes above a specific containment threshold and mapping the reads back to the query genomes to evaluate both mapping detection and effective coverage. In all but one case, k-mer-based genome detection of the query was lower than mapping-based detection - in some cases significantly so. This was also seen with a a smaller set of samples in Irber et al., 2022 [14], and is likely because mapping-based approaches can tolerate mismatches, while k-mer based approaches require exact mismatches.

Second, as designed, branchwater cannot robustly detect sequence similarity for data sets under 10kb in size. This limit is related to the scaling approach used to reduce the effective search space and enable petabase-scale search. Branchwater leverages FracMinHash sketching to build a reduced representation of each data set, which enables direct and accurate sequence similarity and containment comparisons without needing to access the original sequencing reads. Because only a fraction of the original data needs to be stored, FracMinHash sketches are good basic components in the implementation of systems that allow searching large collections of datasets. However, this approach comes with some detection limitations. Robust detection requires a minimum overlap of 2-3 hashes [14]. With the scaling used in branchwater, this represents approximately 2-3 kb of matching sequence. Since many plasmids and most bacterial and archaeal genomes are far larger than 10kb, branchwater is well suited to detecting matches to such sequences in the SRA. Future storage and search optimizations may enable higher resolution search, but for now, sequences smaller than 10kb may be missed as a result of this fractional k-mer selection.

### Tackling biological use cases with branchwater

K-mer search via branchwater has been used in two projects so far - Lumian et al. (2022) [16] and Viehweger et al. (2021) [15]. Viehweger et al. used branchwater to find a metagenome containing an additional *Klebsiella pneumoniae* for a large scale analysis of outbreak data, while Lumian et al. conducted a global biogeography analysis of five new antarctic cyanobacteria. Both studies benefited from the low cost and comprehensive nature of the search.

We expect a broader range and more elaborate set of use cases to emerge as petabase scale search becomes more widely available. The low cost of search with branchwater is particularly enabling for exploratory efforts, although the sheer size of the underlying data needed even for branchwater continues to present obstacles.

There are several scientific limitations of our approach to overcome as well. The current search approach has limited sensitivity to divergent sequence beyond the genus level, and cannot find smaller matches. These are topics for future research and development.

### Following up on Branchwater results

Many Branchwater use cases are intended for early-stage hypothesis generation and refinement, i.e. branchwater implements the first part of a “hit to lead” pipeline. Hence Branchwater operates at an early stage in conceptual and concrete workflows. The initial steps immediately after executing Branchwater are (1) choosing a threshold at which to filter results, (2) evaluating the overall results by type of metagenome retrieved, and (3) retrieving the data underlying the matches.

As branchwater search is exhaustive, the first analysis step taken is typically picking a more stringent threshold. The number of metagenomes results is highly dependent on the query, ranging from dozens of results for organisms from understudied environments to tens or hundreds of thousands of results for organisms from well-sequenced environments, such as human gut search in (3). As a result, thresholds are typically chosen based on the use case and the observed distribution of the annotated metagenome type (`Scienti ficName` from the SRA Runinfo database). After filtering, many paths can be taken. A plethora of general purpose bioinformatics tools exist for working with the data from individual metagenomes.

We have built two custom tools in concert with sourmash and branchwater, genome-grist and spacegraphcats. Genome-grist performs an entirely automated reference-based characterization of individual metagenomes that follows the minimum metagenome cover produced by sourmash gather with mapping of short reads; it is described in Irber et al. [14] and was used in Lumian et al. [16]. While genome-grist does download the entire data set in order to map the reads, it is still reference based and thus relatively lightweight.

spacegraphcats is an assembly-graph based investigative tool for metagenomes that retrieves graph neighborhoods from metagenome assembly graphs for the purpose of investigating strain variation [24]. It was used to retrieve putative accessory elements from sourmash matches in Reiter et al. (2022) [25] and Lumian et al. (2022) [26]. It is much heavier weight than genome-grist because it relies on a compact De Bruijn graph, which is expensive to build for very rich or diverse metagenomes.

### Design alternatives to the current branchwater implementation

The current branchwater software is a simple and effective implementation that is easy to analyze algorithmically and supports a number of use cases. However, many improvements are possible: FracMinHash analyses are based on comparing collections of 64-bit integers, and there are many effective tools and approaches for organizing and searching such collections more efficiently than is presently done.

One area for particular improvement is storing and loading sketches more efficiently. The current JSON-based format is convenient for debugging and multi-language interoperability but is extremely inefficient. Moreover, each file currently contains three k-mer sketches (one per each desired k-mer search size), which means approximately 3 times as much data is loaded per query than is actually used. FracMinHash also could support fractional loading, i.e. decreased resolution by loading only the bottom portion of the sketch; this would enable must faster searches albeit at lower resolution. This is not yet supported by the underlying sourmash library.

Currently the data files are organized as flat files in a single directory on a single network flle system. There are a variety of practical ways to speed up the search by distributing sketch files across multiple nodes, but this is logistically challenging. In particular, our current usage involves running branchwater once every few weeks on our HPC, which does not have sufficient local storage to distribute the data sets across nodes. In addition, the speed savings from distributing sketches across nodes is unlikely to be rewarding enough to offset the maintenance requirements for a distributed collection of 7.5 TB of sketches. Future work could include implementation of an automated distribution system, although careful evaluation of the maintenance and update requirements would be needed.

We could also create a simple pre-filter for each flle using a data structure with one-sided error. For example, we could create a Bloom Alter for each sketch that could be used to estimate containment prior to loading the full sketch flle. However, for some potentially common use cases such as queries with many matches, this could add significant I/O without speeding up the actual search.

Building an inverted index that maps hashes to data sets could also enable rapid queries. Two challenges here are the scale of the catalog and the number of data sets; the total number of hashes present in our metagenome catalog is 375 billion hashes, across nearly 800,000 data sets.

Despite these many opportunities for optimization, we argue that there is a significant benefit to the simplicity of our current approach. In particular, providing the sketches in individual files organized by accession makes it straightforward to access individual sketches by a distinct ID and quickly update the overall metagenome catalog. This is particularly valuable since the sourmash Python package provides a flexible suite of tools for inspecting and manipulating individual metagenome sketches. The lack of auxiliary data structures also avoids expensive load and synchronization steps when adding new datasets. These features are important for downstream user investigation as well as maintainability and correctness, which are important considerations in any scientific software workflow.

## Conclusion

We provide a flexible and fast petabase-scale search based on FracMinHash, together with some simple downstream summarization tools and an increasingly mature (but much slower) investigative ecosystem. This supports and enables a wide range of biological use cases that take advantage of public data; these use cases range from biomedical to ecological to technical (Table 1).

## Data availability statement

All of the original data underlying the Branchwater database is available from the NCBI Sequence Read Archive. A current catalog of the SRA accessions is available at XXX. The sketch collection is 7.5 TB and is available upon request. All sourmash sketches are provided under Creative Commons Zero (CC0) - No Rights Reserved.

## References

1. The Sequence Read Archive R Leinonen, H Sugawara, M Shumway Nucleic Acids Research (2010-11-09) https://doi.org/c652z5 DOI: 10.1093/nar/gkq1019 · PMID: 21062823 · PMCID: PMC3013647

2. A unified catalog of 204,938 reference genomes from the human gut microbiome Alexandre Almeida, Stephen Nayfach, Miguel Boland, Francesco Strozzi, Martin Beracochea, Zhou Jason Shi, Katherine S Pollard, Ekaterina Sakharova, Donovan H Parks, Philip Hugenholtz, … Robert D Finn Nature Biotechnology (2020-07-20) https://doi.org/gg5hgn DOI: 10.1038/s41587-020-0603-3 · PMID: 32690973 · PMCID: PMC7801254

3. A new view of the tree of life Laura A Hug, Brett J Baker, Karthik Anantharaman, Christopher T Brown, Alexander J Probst, Cindy J Castelle, Cristina N Butterfield, Alex W Hernsdorf, Yuki Amano, Kotaro Ise, … Jillian F Banfield Nature Microbiology (2016-04-11) https://doi.org/bpkh DOI: 10.1038/nmicrobiol.2016.48 · PMID: 27572647

4. Clades of huge phages from across Earth’s ecosystems Basem Al-Shayeb, Rohan Sachdeva, Lin-Xing Chen, Fred Ward, Patrick Munk, Audra Devoto, Cindy J Castelle, Matthew R Olm, Keith Bouma-Gregson, Yuki Amano, … Jillian F Banfield Nature (2020-02-12) https://doi.org/ggkq9d DOI: 10.1038/s41586-020-2007-4 · PMID: 32051592 · PMCID: PMC7162821

5. Use of simulated data sets to evaluate the fidelity of metagenomic processing methods Konstantinos Mavromatis, Natalia Ivanova, Kerrie Barry, Harris Shapiro, Eugene Goltsman, Alice C McHardy, Isidore Rigoutsos, Asaf Salamov, Frank Korzeniewski, Miriam Land, … Nikos C Kyrpides Nature Methods (2007-04-29) https://doi.org/b28wj2 DOI: 10.1038/nmeth1043 · PMID: 17468765

6. Evaluating Metagenome Assembly on a Simple Defined Community with Many Strain Variants Sherine Awad, Luiz Irber, CTitus Brown Cold Spring Harbor Laboratory (2017-06-25) https://doi.org/ghvn6x DOI: 10.1101/155358

7. Fast search of thousands of short-read sequencing experiments Brad Solomon, Carl Kingsford Nature Biotechnology (2016-02-08) https://doi.org/f8ddk3 DOI: 10.1038/nbt.3442 · PMID: 26854477 · PMCID: PMC4804353

8. Ultrafast search of all deposited bacterial and viral genomic data Phelim Bradley, Henk C den Bakker, Eduardo PC Rocha, Gil McVean, Zamin Iqbal Nature Biotechnology (2019-02) https://doi.org/gf4x49 DOI: 10.1038/s41587-018-0010-1 · PMID: 30718882 · PMCID: PMC6420049

9. Improved Search of Large Transcriptomic Sequencing Databases Using Split Sequence Bloom Trees Brad Solomon, Carl Kingsford Journal of Computational Biology (2018-07) https://doi.org/gdxhgz DOI: 10.1089/cmb.2017.0265 · PMID: 29641248 · PMCID: PMC6067102

10. Petabase-scale sequence alignment catalyses viral discovery Robert C Edgar, Jeff Taylor, Victor Lin, Tomer Altman, Pierre Barbera, Dmitry Meleshko, Dan Lohr, Gherman Novakovsky, Benjamin Buchfink, Basem Al-Shayeb, … Artem Babaian Nature (2022-01-26) https://doi.org/gn9xsd DOI: 10.1038/s41586-021-04332-2 · PMID: 35082445

11. Home https://www.searchsra.org/

12. MetaGraph: Indexing and Analysing Nucleotide Archives at Petabase-scale Mikhail Karasikov, Harun Mustafa, Daniel Danciu, Marc Zimmermann, Christopher Barber, Gunnar Rätsch, André Kahles Cold Spring Harbor Laboratory (2020-10-02) https://doi.org/ghfd79 DOI: 10.1101/2020.10.01.322164

13. sourmash: a library for MinHash sketching of DNA C Titus Brown, Luiz Irber The Journal of Open Source Software (2016-09-14) https://doi.org/ghdrk5 DOI: 10.21105/joss.00027

14. Lightweight compositional analysis of metagenomes with FracMinHash and minimum metagenome covers Luiz Irber, Phillip T Brooks, Taylor Reiter, NTessa Pierce-Ward, Mahmudur Rahman Hera, David Koslicki, CTitus Brown Cold Spring Harbor Laboratory (2022-01-12) https://doi.org/gn34zt DOI: 10.1101/2022.01.11.475838

15. Context-aware genomic surveillance reveals hidden transmission of a carbapenemase-producing Klebsiella pneumoniae Adrian Viehweger, Christian Blumenscheit, Norman Lippmann, Kelly L Wyres, Christian Brandt, Jörg B Hans, Martin Hölzer, Luiz Irber, Sören Gatermann, Christoph Lübbert, … Brigitte König Microbial Genomics (2021-12-16) https://doi.org/gq5gt8 DOI: 10.1099/mgen.0.000741 · PMID: 34913861 · PMCID: PMC8767333

16. Biogeographic Distribution of Five Antarctic Cyanobacteria Using Large-Scale k-mer Searching with sourmash branchwater Jessica Lumian, Dawn Sumner, Christen Grettenberger, Anne D Jungblut, Luiz Irber, NTessa Pierce-Ward, CTitus Brown Cold Spring Harbor Laboratory (2022-10-30) https://doi.org/gq5p6v DOI: 10.1101/2022.10.27.514113

17. Mash: fast genome and metagenome distance estimation using MinHash Brian D Ondov, Todd J Treangen, Páll Melsted, Adam B Mallonee, Nicholas H Bergman, Sergey Koren, Adam M Phillippy Genome Biology (2016-06-20) https://doi.org/gfx74q DOI: 10.1186/s13059-016-0997-x · PMID: 27323842 · PMCID: PMC4915045

18. Dashing: fast and accurate genomic distances with HyperLogLog Daniel N Baker, Ben Langmead Genome Biology (2019-12) https://doi.org/ggkmjc DOI: 10.1186/s13059-019-1875-0 · PMID: 31801633 · PMCID: PMC6892282

19. Mash Screen: high-throughput sequence containment estimation for genome discovery Brian D Ondov, Gabriel J Starrett, Anna Sappington, Aleksandra Kostic, Sergey Koren, Christopher B Buck, Adam M Phillippy Genome Biology (2019-11-05) https://doi.org/ghtqmb DOI: 10.1186/s13059-019-1841-x · PMID: 31690338 · PMCID: PMC6833257

20. IMPROVING MIN HASH VIA THE CONTAINMENT INDEX WITH APPLICATIONS TO METAGENOMIC ANALYSIS David Koslicki, Hooman Zabeti Cold Spring Harbor Laboratory (2017-09-04) https://doi.org/ghvn6z DOI: 10.1101/184150

21. Large-scale sequence comparisons with sourmash NTessa Pierce, Luiz Irber, Taylor Reiter, Phillip Brooks, CTitus Brown F1000Research (2019-07-04) https://doi.org/gf9v84 DOI: 10.12688/f1000research.19675.1 · PMID: 31508216 · PMCID: PMC6720031

22. Debiasing FracMinHash and deriving confidence intervals for mutation rates across a wide range of evolutionary distances Mahmudur Rahman Hera, NTessa Pierce-Ward, David Koslicki Cold Spring Harbor Laboratory (2022-01-12) https://doi.org/gn342h DOI: 10.1101/2022.01.11.475870

23. Sustainable data analysis with Snakemake Felix Mölder, Kim Philipp Jablonski, Brice Letcher, Michael B Hall, Christopher H Tomkins-Tinch, Vanessa Sochat, Jan Forster, Soohyun Lee, Sven O Twardziok, Alexander Kanitz, … Johannes Köster F1000Research (2021-04-19) https://doi.org/gj76rq DOI: 10.12688/f1000research.29032.2 · PMID: 34035898 · PMCID: PMC8114187

24. Exploring neighborhoods in large metagenome assembly graphs using spacegraphcats reveals hidden sequence diversity CTitus Brown, Dominik Moritz, Michael P O’Brien, Felix Reidl, Taylor Reiter, Blair D Sullivan Genome Biology (2020-07-06) https://doi.org/d4bb DOI: 10.1186/s13059-020-02066-4 · PMID: 32631445 · PMCID: PMC7336657

25. Meta-analysis of metagenomes via machine learning and assembly graphs reveals strain switches in Crohn’s disease Taylor E Reiter, Luiz Irber, Alicia A Gingrich, Dylan Haynes, NTessa Pierce-Ward, Phillip T Brooks, Yosuke Mizutani, Dominik Moritz, Felix Reidl, Amy D Willis, … CTitus Brown Cold Spring Harbor Laboratory (2022-07-05) https://doi.org/gq5gt9 DOI: 10.1101/2022.06.30.498290

26. Metabolic Capacity of the Antarctic Cyanobacterium Phormidium pseudopriestleyi That Sustains Oxygenic Photosynthesis in the Presence of Hydrogen Sulfide Jessica E Lumian, Anne D Jungblut, Megan L Dillion, Ian Hawes, Peter T Doran, Tyler J Mackey, Gregory J Dick, Christen L Grettenberger, Dawn Y Sumner Genes (2021-03-16) https://doi.org/gjmxx5 DOI: 10.3390/genes12030426 · PMID: 33809699 · PMCID: PMC8002359

